# Reduced fronto-striatal volume in ADHD in two cohorts across the lifespan

**DOI:** 10.1101/790204

**Authors:** Renata Basso Cupertino, Sourena Soheili-Nezhad, Eugenio Horacio Grevet, Cibele Edom Bandeira, Felipe Almeida Picon, Maria Eduarda Tavares, Jilly Naaijen, Daan van Rooij, Sophie Akkermans, Eduardo Schneider Vitola, Marcel P Zwiers, Pieter J. Hoekstra, Vitor Breda, Jaap Oosterlaan, Catharina A Hartman, Christian F. Beckmann, Jan K. Buitelaar, Barbara Franke, Claiton Henrique Dotto Bau, Emma Sprooten

## Abstract

**Objective:** Neuroimaging studies have associated Attention-Deficit/Hyperactivity Disorder (ADHD) with altered brain anatomy. However, small and heterogeneous study samples, and the use of region-of-interest and tissue-specific analyses have limited the consistency and replicability of these effects. The present study uses a fully data-driven multivariate approach to investigate alterations in both gray and white matter simultaneously, and capture neuroanatomical features associated with ADHD in two large, independent, demographically different cohorts.

**Methods:** The study comprised two ADHD cohorts with structural magnetic resonance imaging data: the Dutch NeuroIMAGE cohort (n=890, average age 17.2 years, discovery sample) and the Brazilian IMpACT cohort (n=180, average age 44.2 years, cross validation sample). Using independent component analysis of whole-brain morphometry images in the NeuroIMAGE cohort, 375 independent components of neuroanatomical variations were extracted and assessed their association with ADHD. Afterwards, ADHD-associated components were cross validated in the Brazilian IMpACT cohort.

**Results:** In both discovery (corrected- *p*=0.020) and validation (*p=*0.033) cohorts, ADHD diagnosis was significantly associated with reduced brain volume in a component mapping to frontal lobes, striatum, and their interconnecting white-matter tracts. The most pronounced case-control differences were localized in white matter adjacent to the orbitofrontal cortex.

**Conclusion:** Independent component analysis is a sensitive approach to uncover neuroanatomical alterations in ADHD and avoid bias attributable to *a priori* region-of-interest based methods. Current results provide further evidence for the role of the fronto-striatal circuit in ADHD. The fact that the two cohorts are from different continents and comprising different age ranges highlights the robustness of the findings.

## Introduction

Attention-Deficit Hyperactivity Disorder (ADHD) is one of the most common psychiatric disorders worldwide, characterized by age-inappropriate levels of inattention and/or hyperactivity/impulsivity leading to significant impairment. Its prevalence is about 5% in children and about 3% in adults (1,2). ADHD is clinically highly heterogeneous, and around 70% of the affected individuals present with comorbid psychiatric disorders, further complicating diagnosis and treatment (3,4). The clinical profile of ADHD changes throughout development: while children are more likely to present with symptoms of hyperactivity/impulsivity, adolescents and adults often experience more symptoms of inattention (1,5). Cognitive functions such as inhibitory control and working memory (6), emotion regulation (7), and motivational processes (8) are also affected in many individuals with ADHD across the lifespan.

Although the underlying neurobiology of ADHD is only partly understood, neuroimaging studies have identified several structural and functional brain changes associated with this disorder (9–12). The most consistent findings in structural magnetic resonance imaging (MRI) studies on ADHD point to a reduction in total brain volume and grey matter in individuals with ADHD compared to controls (10,11,13,14). More specifically, Sáenz et al (10) associated ADHD with structural alterations in the basal ganglia, prefrontal cortex, and the corpus callosum. A mega-analysis comprising over 3,000 individuals found smaller volumes of five out of seven subcortical structures in ADHD, with strongest effects observed in the amygdala, nucleus accumbens, and putamen (13). Age-stratified analyses in that study showed that the effect of ADHD on brain structure was stronger in children, and no statistically significant effect was seen in adults above the age of 22 years (13). More recently, a coordinated analysis involving 36 centers showed that children with ADHD also have reduced cortical surface area, especially in frontal, cingulate, and temporal regions as well as reduced cortical thickness in fusiform gyrus and temporal pole (15). That study also did not find differences in surface area and cortical thickness in the adolescent and adult groups, again suggesting an age-dependent effect. Evidence from studies using diffusion-weighted imaging implicates white-matter microstructural alterations in ADHD (16–18), possibly hampering neural communication amongst and between cortical and subcortical areas. Two meta-analyses found altered fractional anisotropy in widespread regions in patients with ADHD, with the most consistent findings in the corpus callosum, anterior corona radiata, right forceps minor, bilateral internal capsule, and left cerebellum (9,17).

Previous structural MRI studies in ADHD either focused on *a priori*, automatically segmented, cortical or subcortical areas delimiting specific regions of interest (ROIs) (11,13), or explored changes in brain volume and cortical area and thickness using mass univariate methods at voxel or vertex resolution (19–21). While these methods are primarily sensitive to the detection of grey-matter differences, evidence from diffusion-weighted imaging shows that ADHD may also affect white matter microstructure (9,16,18). In the present study, we used a combination of tensor-based morphometry (TBM) and independent component analysis (ICA), allowing us to optimize sensitivity to the detection of local differences in both grey- and white-matter tissue and their spatial covariation, in a whole-brain multivariate analysis.

ICA has been frequently used in functional MRI studies, leading to the identification of several functional brain networks (22,23). More recently, ICA has also been successfully applied to whole-brain diffusion imaging data (24,25). In relation to ADHD, multi-modal linked ICA has been used for data fusion across different MRI modalities (cortical thickness and area, voxel-based morphometry, and diffusion tensor imaging), showing association of ADHD symptom severity with several independent components of grey and white matter properties distributed across many brain regions (26–28). However, replication in neuroimaging remains scarce, and the generalizability across populations of different ethnic, social, and cultural backgrounds is unknown. Furthermore, it remains unclear how any potential neuroimaging markers of ADHD, being a neurodevelopmental disorder, vary with age across the lifespan.

The present study aimed to identify structural brain differences in association with ADHD across the lifespan in two relatively large, independent cohorts. Using TBM, we created images capturing local brain volume variation in both grey and white-matter. Subsequently, ICA was used to isolate independent components of spatial covariance from these images. We assessed association of the independent brain components with ADHD diagnosis and symptom dimensions in a discovery sample of adolescent and young adults (NeuroIMAGE). Afterwards, we appraised the effect of a significantly associated component in an independent and clinically different validation sample of individuals diagnosed with ADHD during adulthood (Brazilian IMpACT cohort).

## Material and Methods

### Samples

#### NeuroIMAGE cohort

The NeuroIMAGE cohort is a Dutch prospective multi-site study aimed to investigate the longitudinal course of ADHD relying on two MRI waves (NeuroIMAGE I/II). Details of this cohort have been described elsewhere (29). Participants were enrolled at two sites, the Vrije Universiteit in Amsterdam and the Radboud University Medical Center in Nijmegen. The study was approved by the local ethics committees, and written informed consent was obtained from all participants and their legal guardians. The cohort included unrelated participants (n=138) as well as those with full sibling relationships in families of different sizes (n=241 sibling pairs, n=70 three siblings, n=15 four siblings). In order to maximize sample size, data from the two longitudinal study waves were combined such that individuals who participated twice were only included at the first wave (NeuroIMAGE I). The ADHD diagnosis was primarily based on a semi-structured clinical interview using a Dutch translation of the Schedule for Affective disorders and Schizophrenia for school-age children – present and lifetime version (K-SADS-PL), based on the fourth version of the diagnostic and statistical manual (DSM-IV). To optimize the diagnostic assessment, the information from the K-SADS-PL interview was combined with information from the Conners Adult ADHD Rating Scale (CAARS R-L), Conners Parent Rating Scale (CPRS R:L), and Conners Wells Adolescent Self-Report Scale: Short Form (CASS:S). For an ADHD full diagnosis, individuals had to have six or more symptoms in the inattention domain and/or in the hyperactivity/impulsivity domain causing impairment in multiple settings, as well as a Conners T-score ≥63. Unaffected individuals had ≤3 symptoms *and* a Conners T-score <63. Participants who did not fulfill criteria for either category were classified as *subthreshold* ADHD. Psychiatric comorbidities, such as anxiety, depression, and oppositional behavior were assessed by the Dutch version of the Strengths and Difficulties Questionnaire (SDQ). Exclusion criteria were: Intelligence Quotient (IQ) <70, diagnosis of autism, neurological disorders, such as epilepsy, general learning difficulties, brain trauma, or known genetic disorders, such as Fragile X or Down syndrome.

From the NeuroIMAGE I study wave, 807 individuals were enrolled for a scan session. In both sites, 1.5T MRI scanners were employed (Siemens Magnetom SONATA and AVANTO, Erlangen Germany), using 8-channel phased array head coils. T_1_-weighted anatomical scans were acquired at an isotropic resolution of 1mm using a 3D magnetization prepared rapid acquisition with gradient echoes (MPRAGE) sequence with 176 slices, flip angle=7°, TE=2.95ms, TR=2,730ms, TI=1,000ms, matrix size=256×256, and parallel acquisition (GRAPPA) with an acceleration factor of 2. From NeuroIMAGE II study wave, 87 subjects were considered, only from one site (Nijmegen); MRI scanner and image acquisition parameters remained the same as the first wave. After quality control, structural brain MRI scans of a total of 890 individuals were considered (359 affected, 98 subthreshold, and 433 unaffected; ages range from 7 to 29 years). Sample characteristics of this cohort are provided in **Table 1**.

**Table 1.** Demographic and clinical characteristics of the NeuroIMAGE cohort.

|  | ADHD<br>(N=359) |  | Subthreshold<br>(N=98) |  | Controls<br>(N=433) |  |
| --- | --- | --- | --- | --- | --- | --- |
|  | *n* | % | *n* | % | *n* | % |
| Sex (male) | 247 | 68.8 | 56 | 57.1 | 206 | 47.6 |
| Scan site (Nijmegen) | 195 | 54.3 | 48 | 49.0 | 203 | 46.9 |
| Wave drawn (NeuroIMAGE I) | 336 | 93.6 | 85 | 86.7 | 386 | 89.1 |
| Age (years)<sup>a</sup> | 17.0 | 3.7 | 18.1 | 3.9 | 17.2 | 3.8 |
| *Comorbidities (lifetime)* |  |  |  |  |  |  |
| Oppositional Defiant Disorder | 109 | 30.4 | 7 | 7.1 | 7 | 1.6 |
| Conduct disorder | 22 | 6.1 | 2 | 2.0 | 0 | 0 |
| Major depressive disorder | 4 | 1.1 | 0 | 0 | 2 | 0.5 |
| Generalized anxiety disorder | 6 | 1.7 | 2 | 2.0 | 2 | 0.5 |
| Avoidant/social phobia disorder | 3 | 0.8 | 0 | 0 | 1 | 0.2 |
| Panic disorder | 0 | 0 | 1 | 1.0 | 0 | 0 |
| *Symptoms (KSADS)<sup>a</sup>* |  |  |  |  |  |  |
| Inattention | 6.6 | 1.9 | 3.6 | 1.4 | 1.2 | 0.9 |
| Hyperactivity/Impulsivity | 5.2 | 2.4 | 2.8 | 1.5 | 1.4 | 1.0 |

<sup>a</sup>Data represented as mean (standard deviation).

#### Brazilian IMpACT cohort

The Brazilian IMpACT cohort was assessed by the ADHD Outpatient Program – Adult Division at the Hospital de Clínicas de Porto Alegre (PRODAH-A). Further information of this cohort have been described elsewhere (30,31). All participants were unrelated adults of white (European descent) Brazilian ethnicity aged 18 years or older. ADHD diagnosis was based on DSM-5 diagnostic criteria, using the K-SADS epidemiologic version (KSADS-E) adapted for adults. Individuals were recruited when seeking for psychiatric help (cases) or donating blood (controls). The Structured Clinical Interview for DSM-IV Axis I Disorders (SCID-I) was used for assessing other lifetime psychiatric comorbidities. Individuals with significant neurological disease, head trauma, history of psychosis, and/or an estimated IQ score below 70 were excluded. Participants signed an informed consent form before the study, which was approved by the Ethics Committee of the hospital. Sample characteristic can be found in **Table 2**.

**Table 2.** Demographic and clinical characteristics of the Brazilian IMpACT cohort.

|  | ADHD<br>(N=118) |  | Controls<br>(N=62) |  |
| --- | --- | --- | --- | --- |
|  | *n* | % | *n* | % |
| Sex (male) | 51 | 43.2 | 33 | 53.2 |
| Age (years)<sup>a</sup> | 46.9 | 10.5 | 39.2 | 9.6 |
| *Comorbidities (lifetime)* |  |  |  |  |
| Generalized Anxiety Disorder | 77 | 65.3 | 18 | 29.0 |
| Oppositional Defiant Disorder | 82 | 69.5 | 4 | 6.4 |
| Major Depressive Disorder | 53 | 44.9 | 23 | 37.1 |
| Social Phobia | 43 | 36.4 | 11 | 17.8 |
| Bipolar Disorder* | 36 | 30.5 | 3 | 4.8 |
| Substance Use Disorder | 33 | 28.0 | 5 | 8.1 |
| Eating Disorders <sup>c</sup> | 25 | 21.2 | 8 | 12.9 |
| Obsessive Compulsive Disorder | 26 | 22.0 | 2 | 3.2 |
| Antisocial Personality Disorder | 4 | 3.4 | 1 | 1.6 |
| *Symptoms (KSADS) <sup>a</sup>* |  |  |  |  |
| Inattention | 6.3 | 2.5 | 1.3 | 2.3 |
| Hyperactivity/Impulsivity | 4.6 | 2.7 | 1.2 | 1.6 |

<sup>a</sup> Data represented as mean (standard deviation). <sup>b</sup> Including other specified bipolar and related disorder cases. <sup>c</sup> including anorexia nervosa, bulimia nervosa or binge-eating.

Participants were re-evaluated on average 13 years after diagnosis and underwent MRI scanning, using a 3.0T Siemens SPECTRA scanner and a 12-channel head coil. A high-resolution structural MRI volume was acquired using a T_1_-weighted 3D MPRAGE sequence with 192 slices, flip angle=7°, TE=2.55ms, TR=2,530ms, TI=1,100ms, matrix size=256×256, isotropic resolution of 1mm, and a GRAPPA factor of 2. After quality control procedures, 180 individuals with structural MRI scan were included (118 affected and 62 unaffected; age range from 26 to 74 years).

### Imaging preprocessing

All T1-weighted images were corrected for magnetic field bias using the N4 algorithm (32), and the brain field of view was cropped using the FSL standard_space_roi tool. For each cohort separately, images were registered to an average space to create a cohort-specific minimum deformation brain template (**Supplementary Figure 1**). Four iterations of linear registration and five iterations of diffeomorphic SyN registration were used for template creation (33). The nonlinear warps that transformed each subject’s native brain volume to the common template were used to derive Jacobian determinant fields that encode local brain volume variation across the study individuals. The Jacobian values were subsequently log-transformed to symmetrize their distribution around zero to obtain Jacobian determinant maps per participant. The Jacobian values were normalized within a brain mask in the standard template space, hence removing the global brain size effect.

**Figure 1.**
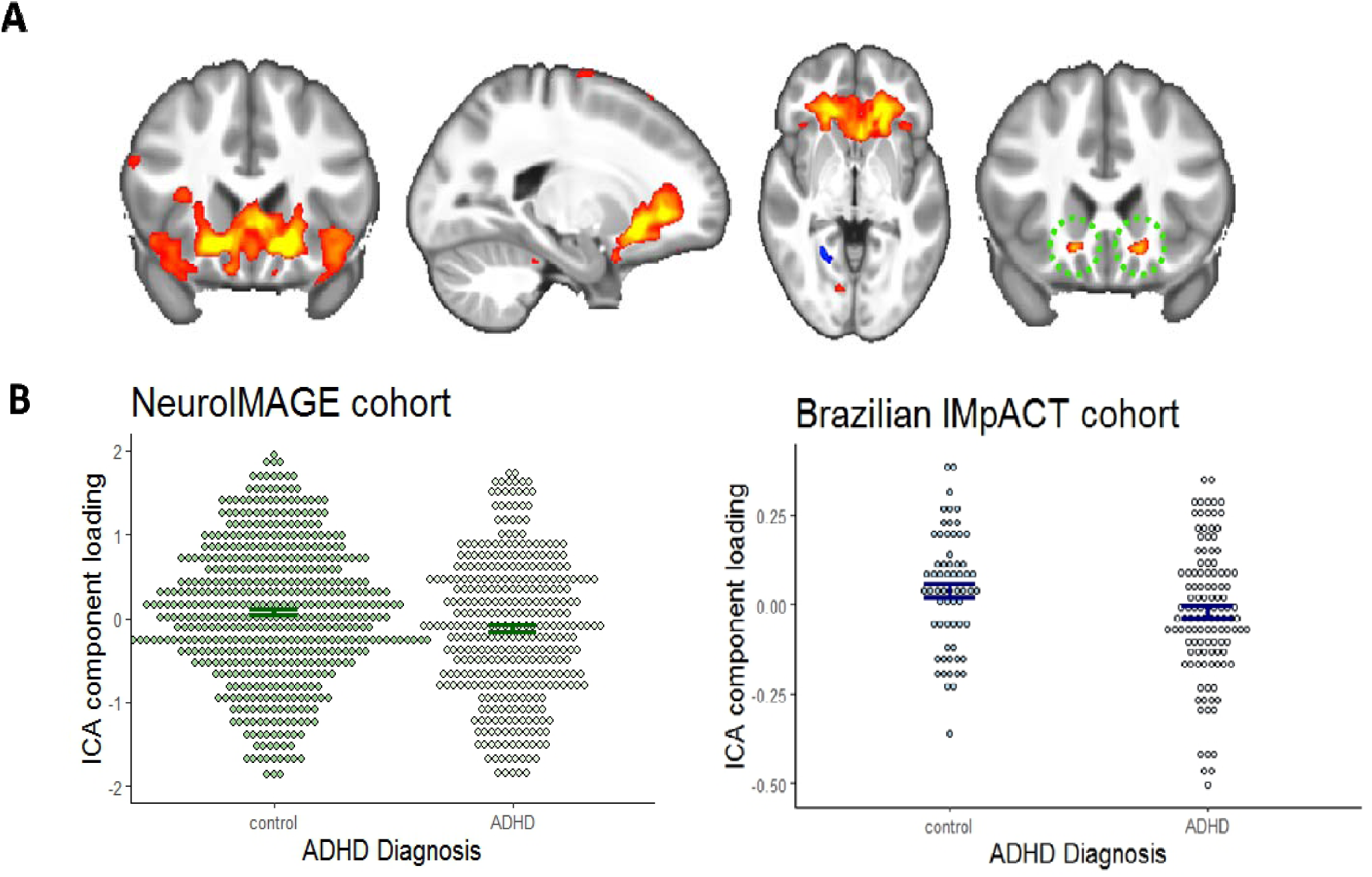
ICA component associated with ADHD. **A.** ADHD patients demonstrated reduced loading of a component capturing volume of prefrontal white-matter together with orbitofrontal, striatal, and insular grey-matter (IC z-score>3.6). The focal area of highest component probability (IC z-score>8) is depicted on the right, showing orbitofrontal white-matter volume reduction bilaterally. **B.** Residualized ICA component loading in individuals with and without ADHD in both cohorts. Error bar indicates standard error.

### Analysis

The Jacobian determinant maps of the NeuroIMAGE cohort (n = 890) were decomposed into spatially independent components using MELODIC, a probabilistic ICA method (23). Four decompositions were performed with varying dimensionality (25, 50, 100 and 200 components) to allow the identification of independent spatial sources in the Jacobian fields at various levels of spatial specificity. Thus, a total of 375 components were extracted. ICA performs a linear matrix factorization of the Jacobian determinant maps resulting in 2 matrices: a matrix of spatially independent component maps reflecting the contribution of each voxel to each component; and a mixing matrix containing the loading of each individual on each component. The latter was used to test case-control differences in each spatial component.

In the discovery cohort (NeuroIMAGE), a permutation-based general linear model (PALM) (34) was used to correct for multiple testing of all 375 brain components loading values with ADHD, while controlling for age, sex, scan site and study wave. To control for family structure in the NeuroIMAGE sample, permutation analysis was used with exchangeability blocks that only allow permutations either within unrelated participants or siblings-blocks separately, or whole-block permutations across families of the same sizes (35). Type I error rate was controlled across all tested brain components using 10,000 random permutations.

To test the significant NeuroIMAGE-based ICA component in the Brazilian IMpACT cohort, the brain component significantly associated with ADHD in the discovery cohort was mapped to the brain template of the Brazilian IMpACT cohort using non-linear SyN transformation. Spatial regression was used to derive the level of brain volume in this spatial component in each subject of the Brazilian IMpACT cohort. The obtained brain volume parameter, now containing the values of the NeuroIMAGE-based ICA-feature for the subjects in Brazilian IMpACT cohort, was used as an imaging feature and its association with ADHD was assessed using linear regression, controlling for sex and age confounders.

## Results

### NeuroIMAGE cohort

A total of 375 components were decomposed by probabilistic ICA showing areas of structural brain covariation across individuals, of them 52 components were nominally associated (*p*_*uncorrected*_<0.05) with current diagnosis of ADHD. Only one component spanning frontal lobes and striatum remained significant after multiple testing correction. This component showed reduced loading in individuals with current ADHD (n = 359, participants, mixed model *t* = −3.61: uncorrected *p* = 3×10^−4^, permutation-corrected *p* = 0.0085) and lifetime history of ADHD (n = 418 participants, mixed model *t* = −3.19, uncorrected *p* mixed model = 3×10^−4,^ permutation-corrected *p* = 0.0015). The maximal focus of brain volume reduction in this component localized to the bilateral fronto-striatal white-matter adjacent to the orbitofrontal cortex – **Figure 1**). This association remained significant after exclusion of individuals with a positive history of ADHD medication (n = 197 drug naïve patients, *p =* 9×10^−4^; *t* = −3.36).

There was significant correlation of the fronto-striatal component with Conners’ ADHD symptom dimensions; inattention *p =* 0.012, *t* = −2.53; hyperactivity *p =* 0.003, *t* = −3.01), and with the number of hyperactivity/impulsivity symptoms (*p =* 0.040, *t* = −2.06) but not with the number of inattention symptoms (*p =* 0.21; *t* = −1.26) assessed by the KSADS. The fronto-striatal component was correlated with subjects’ sex (reduced in females compared to males; *p* < 0.0001, *t* = 9.35) and age (greater volume reductions with age; *p* < 0.0001, *t* = −5.46), although no interaction effects with sex or age were observed (*p =* 0.92. and 0.27, respectively). The smaller group of participants with subthreshold ADHD also showed similar trend for reduced brain volume in the fronto-striatal component (current subthreshold diagnosis n = 98 subjects, *p =* 0.11, *t* = −1.61; lifetime subthreshold diagnosis n = 74, *p =* 0.13, *t = −1.54*).

### Brazilian IMpACT cohort

The association of the fronto-striatal ICA component was replicated in the Brazilian IMpACT cohort with current ADHD (*p* = 0.032, *t* = −2.16) and with lifetime history of ADHD (*p =* 0.021, *t* = −2.33), also controlling for age and sex (**Table 3** and **Figure 1**). Again, this result remained significant when excluding individuals currently under pharmacological treatment (*p =* 0.030, *t* = −2.18). The same direction of effect was observed as in NeuroIMAGE: adults with ADHD showed smaller brain volume in the fronto-striatal component. The component was also associated with the number of hyperactivity/impulsivity symptoms (*p =* 0.046, *t =* −2.011), where a larger number of symptoms was associated with reduced brain volume. The component was associated with sex (*p* < 0.001, *t* = 5.47), but not with age (*p =* 0.75). There was no interaction of lifetime ADHD with sex (*p =* 0.412) or age (*p =* 0.076) on the component.

**Table 3.**
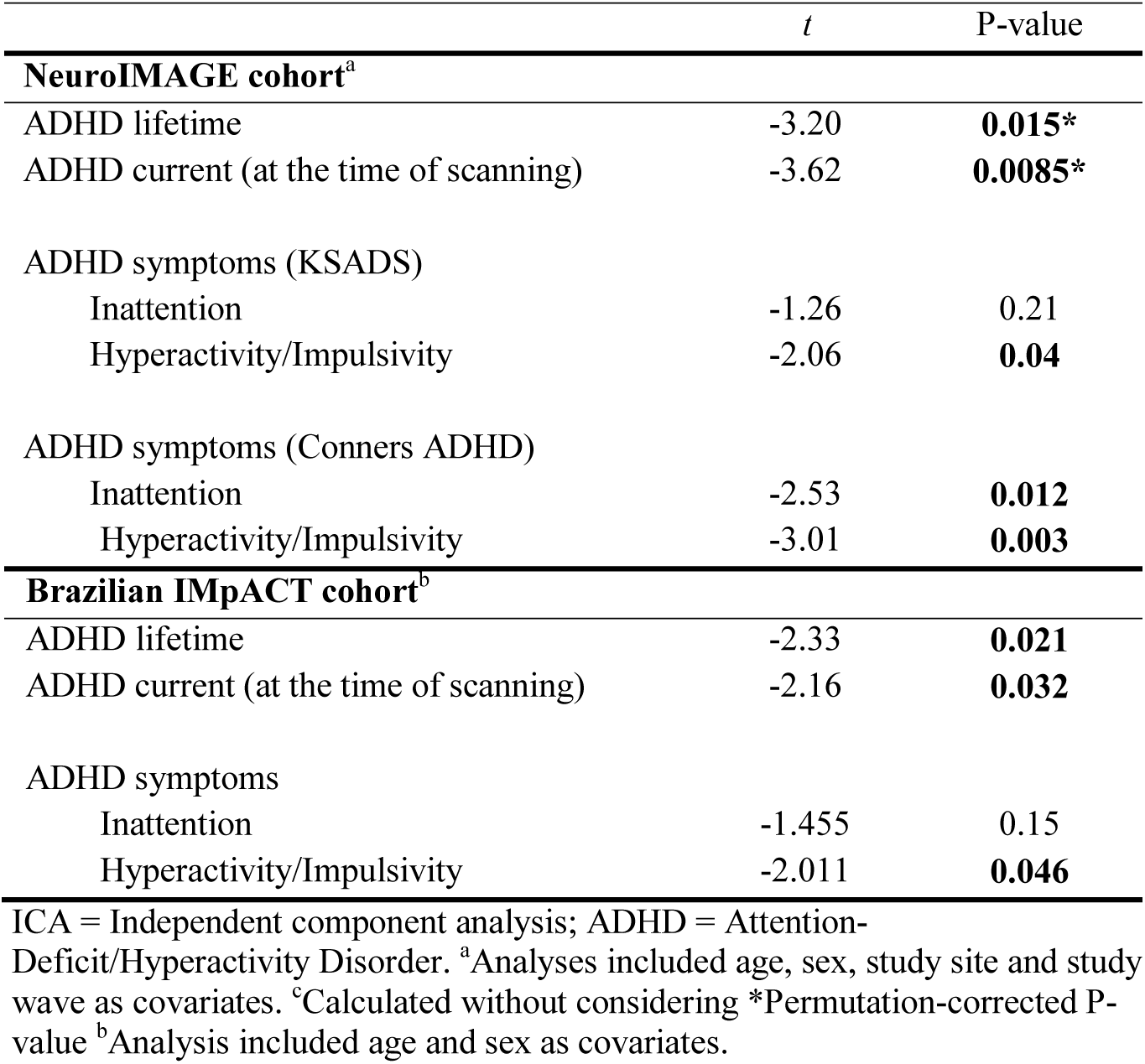
Loading of the ICA component spanning frontal lobes and striatum in both cohorts.

### Secondary Analyses

Considering the broad age-range of the two cohorts, and previously reported significant age-by-diagnosis interactions for brain volumes (36) or differences in brain volumes across the lifespan (13,15), other ICA components were also investigated for possible interactions with age. To reduce the number of tests, only components of the 200-component dimension were investigated. In the NeuroIMAGE cohort, no ICA component showed significant interaction effects after multiple testing correction; however, a nominal effect was observed for 14 components (**Supplementary Figure 2**). The strongest nominal finding was found for a component mapping to bilateral putamen (ICA 17 – **Figure 2**), where participants with ADHD had a slower decrease in regional brain volume with age compared to controls (P_uncorrected_ = 0.0043 – **Figure 2**). The 14 components showing nominal age-by-diagnosis interaction effect in the NeuroIMAGE cohort were also investigated in the Brazilian IMpACT cohort (**Supplementary Figure 2**). In the adult cohort, a nominal interaction effect was observed only for the putamen-related component (ICA 17; P_uncorrected_ = 0.025; **Figure 2**), in the same direction as in the NeuroIMAGE cohort, with the patients showing a less steep rate of decline with age compared to the controls.

**Figure 2.**
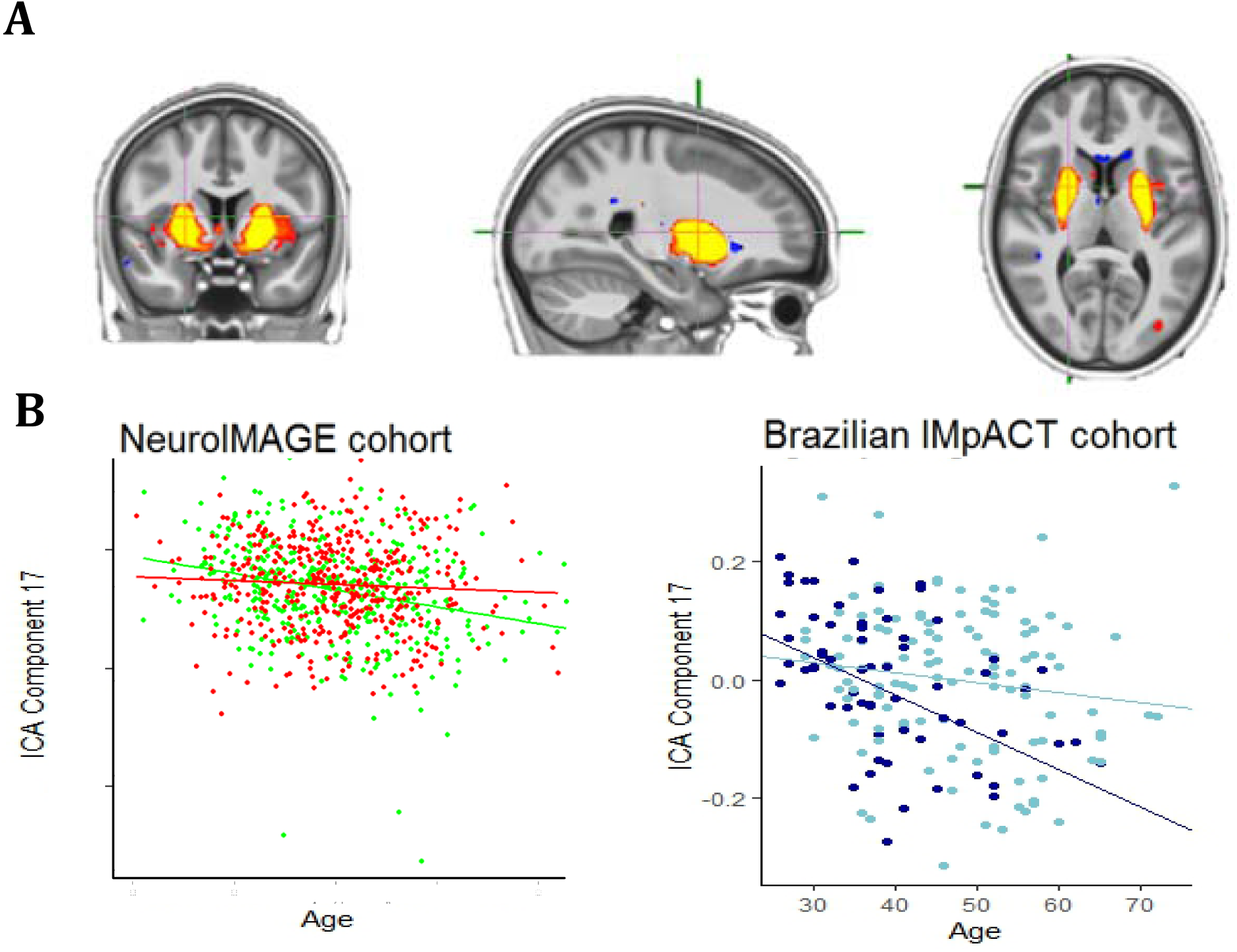
Age-by-diagnosis interaction **A.** ICA component with strongest nominal age-by-diagnosis interaction effect. **B.** Plots of the age-by-diagnosis interaction on ICA component 17 in both cohorts. NeuroIMAGE cohort plots: cases are represented by red dots and dashed lines, and controls by green dots and solid lines. Brazilian IMpACT cohort plots: cases are represented by light blue and controls by dark blue.

## Discussion

This study sought to identify structural brain associations with ADHD using a different approach from those used in most structural MRI studies. The presented data-driven approach was sensitive to both grey- and white-matter volume variation, and the multivariate decomposition into independent spatial sources increased sensitivity to detect whole brain variation that is correlated, both spatially and across individuals. The results provide new evidence for the role of the fronto-striatal circuitry and point to the importance of white-matter in ADHD pathophysiology. Importantly, our work shows the robustness of this finding, as it was seen in two independent cohorts, which were geographically distant and comprised different stages of the lifespan. Also, the present study reported another component with a nominal age-by-diagnosis interaction effect in line with age-dependent effects previously observed in structural changes in ADHD.

A significant case-control brain volume difference was observed for a brain component localized to bilateral fronto-striatal white matter, adjacent to the orbitofrontal cortex. Fronto-striatal circuits are implicated in complex behaviors, such as reward processing, emotion regulation, inhibition, and motivational states (37), and the dysfunction of these circuits is implicated in several psychiatric disorders (37–39), including ADHD (40). Previous structural MRI studies have shown volume reductions in frontal lobes and the striatum (10,13) in individuals with ADHD compared to controls. The present study indicates that this fronto-striatal trait is age-independent across adolescence and adulthood in two independent cohorts of different age ranges and shows for the first time that this fronto-striatal trait alterations generalizes to adults with ADHD up to middle age.

Contrary to most studies that use *a priori* segmented areas, this study used data-driven ICA for separation of raw data into linearly mixed spatial sources. Such decomposition of voxel-wise brain morphometry is useful to discover spatial features that covary beyond *a priori* defined regional boundaries, while avoiding mass univariate voxel wise tests. Extracting morphometric sources without predefined anatomical boundaries assists in deriving MRI features that may better reflect underlying pathophysiological processes. Moreover, using this approach, the maximal focus of brain volume deficits was detected in the bilateral frontal-striatal white-matter, suggesting the importance of considering both grey- and white-matter in conjunction.

Given that several studies, including the largest ADHD neuroimaging mega-analyses to date (13,15), suggested age-dependent associations of ADHD with brain anatomy, we also performed an exploratory analysis to identify potentially age-dependent associations of the ICA components with ADHD. We observed one nominal, but replicated, age-by-diagnosis interaction effect. In line with Hoogman et al. (13) and Greven et al. (36 – also in NeuroIMAGE cohort), this component showing a nominal age-by-diagnosis interaction effect is mapped in the bilateral putamen, where the age-related volume decline occurred at slower rate in ADHD compared to the control group.

It is important to consider this study in the context of some strengths and limitations. This is a cross-sectional study of adolescents and adults, limiting conclusions with regard to brain structural changes across the lifespan and conclusions about developmental aspects, for which longitudinal imaging data would further increase information content. Nevertheless, considering both cohorts, the study includes individuals of a wide age-range, even overlapping ages, allowing us to infer that the main result observed was independent of age.

In conclusion, using tensor-based morphometry driven by both grey- and white-matter, the present findings reinforce the importance of fronto-striatal circuitry and medial frontal white-matter in ADHD neurobiology. Identification of brain structural differences between individuals with and without ADHD provides new insights into the biology underlying this disorder and can contribute to improving diagnosis and treatment in the future. For this, the neural substrate observed here might be of particular interest, given its cross-cultural and age-independent validity.

## Acknowledgments

The NeuroIMAGE study was supported by NIH Grant R01MH62873 (to Stephen V. Faraone), NWO Large Investment Grant 1750102007010 (to Jan Buitelaar), ZonMW grant 60-60600-97- 193, NWO grants 056-13-015 and 433-09-242, and matching grants from Radboud University Nijmegen Medical Center, University Medical Center Groningen and Accare, and Vrije Universiteit Amsterdam. The research leading to these results also received support from the European Community’s Seventh Framework Programme (FP7/2007-2013) under grant agreement number 278948 (TACTICS). Brazilian IMpACT was financed by Conselho Nacional de Desenvolvimento Científico e Tecnológico (Grants 476529/2012-3, 466722/2014-1 and 424041/2016-2), the Coordenação de Aperfeiçoamento de Pessoal de Nível Superior – Brasil (CAPES) – Finance Code 001 and FIPE-HCPA 160600. Barbara Franke was supported by a personal Vici grant from the Innovation Program of the Netherlands Organization for Scientific Research (NWO; grant 016-130-669). Renata B Cupertino was supported by International Brain Research Organization-Latin America (IBRO LARC) Exchange Fellowship. Emma Sprooten is funded by a Hypatia Fellowship (Radboudumc) and a NARSAD Young Investigator Grant (Brain and Behavior Research Foundation, ID: 25034).

This study is part of the International Multicentre persistent ADHD Collaboration (IMpACT; www.impactadhdgenomics.com). IMpACT unites major research centres working on the genetics of ADHD persistence across the lifespan and has participants in the Netherlands, Germany, Spain, Norway, the United Kingdom, the United States, Brazil and Sweden. Principal investigators of IMpACT are: Barbara Franke (chair), Andreas Reif (co-chair), Stephen V. Faraone, Jan Haavik, Bru Cormand, J. Antoni Ramos-Quiroga, Marta Ribases, Philip Asherson, Klaus-Peter Lesch, Jonna Kuntsi, Claiton H.D. Bau, Jan K. Buitelaar, Alejandro Arias Vasquez, Tetyana Zayats, Henrik Larsson, Alysa Doyle, and Eugenio H. Grevet.

