## Supplementary Figure 1 for "Reduced fronto-striatal volume in ADHD in two cohorts across the lifespan"

**Supplementary Figure 1.** Brain templates of both cohorts constructed by iterative registration of all T1-weighted volumes using affine (left) and nonlinear SyN algorithms (middle: first SyN iteration; right: fifth SyN iteration).

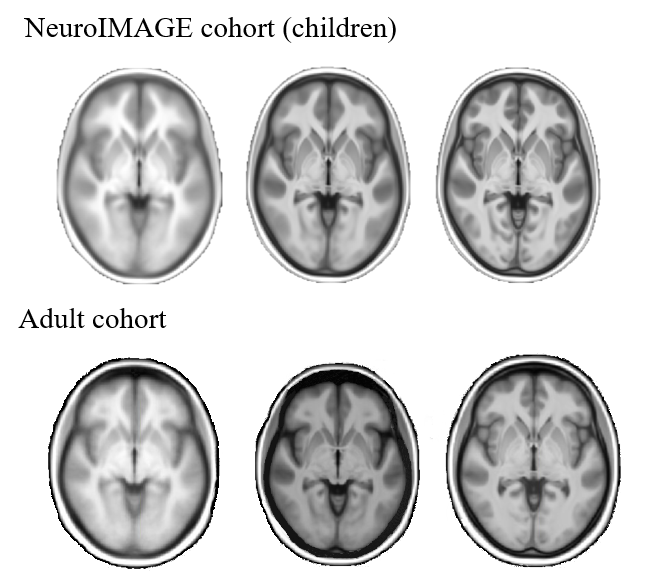

Brazilian IMpACT cohort

NeuroIMAGE cohort

**Supplementary Figure 2.** Fourteen ICA components with nominal age-by-diagnosis interaction effect in the NeuroIMAGE cohort, investigated also in the Brazilian IMpACT cohort. Component 17 (top left), the top component in the discovery sample, was the only component with a significant effect of a consistent direction in both cohorts. NeuroIMAGE cohort plots: cases are represented by red dots and dashed lines, and controls by green dots and solid lines. Brazilian IMpACT cohort plots: cases are represented by light blue and controls by dark blue.

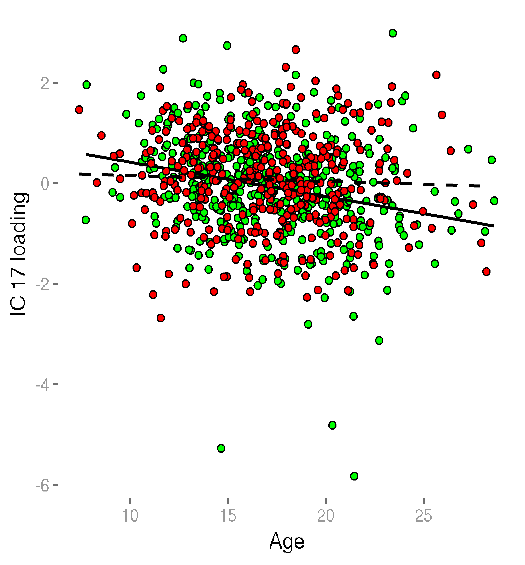

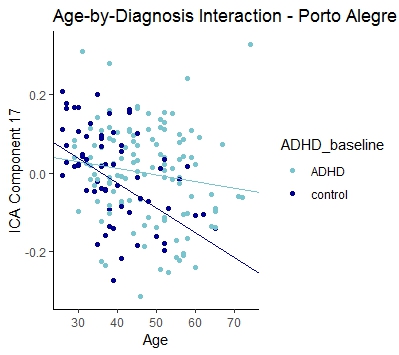

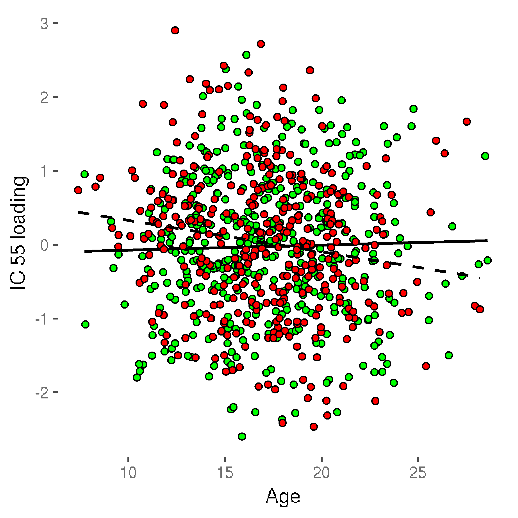

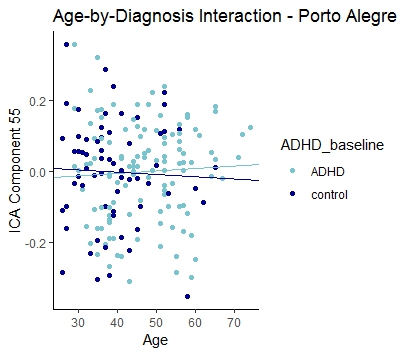

*p =* 0.563

*p =* 0.0043

*p =* 0.0044

*p =* 0.0255

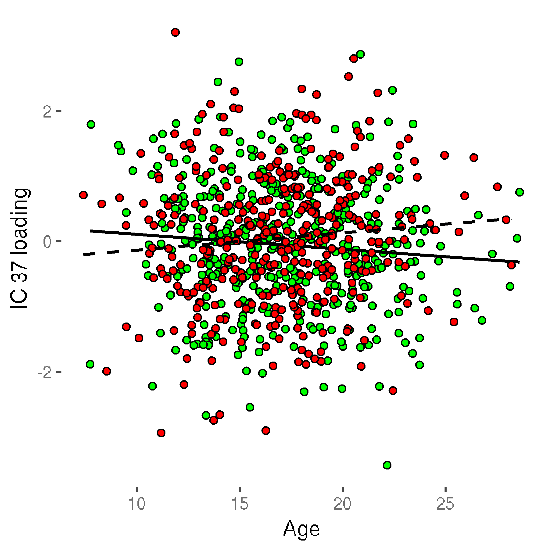

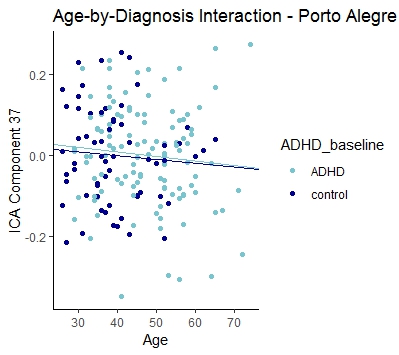

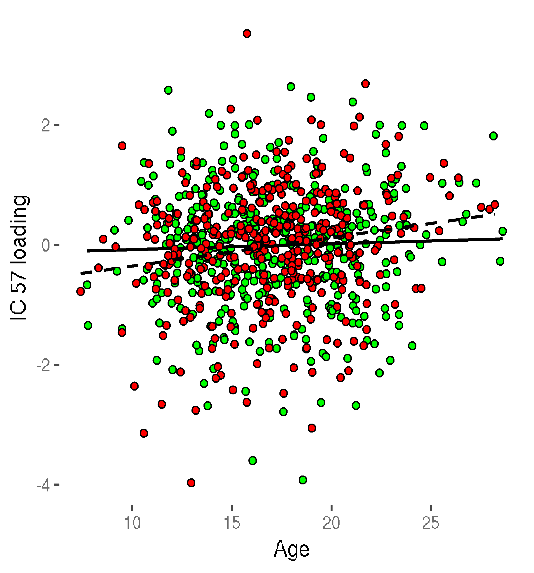

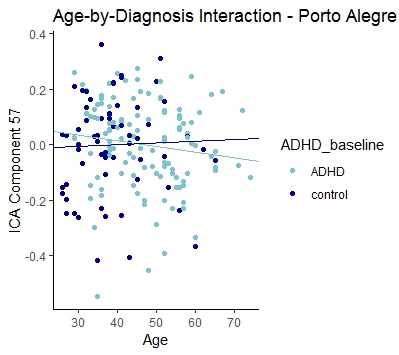

*p =* 0.0161

*p =* 0.3158

*p =* 0.9136

*p =* 0.0058

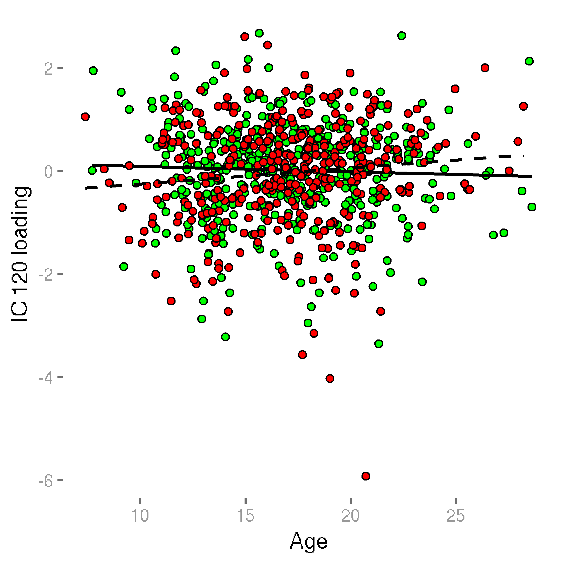

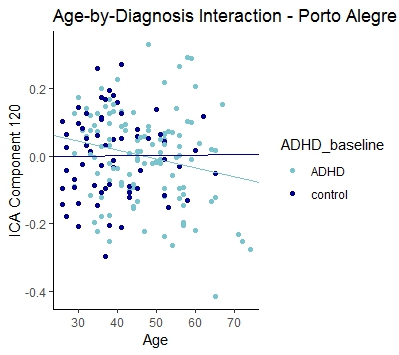

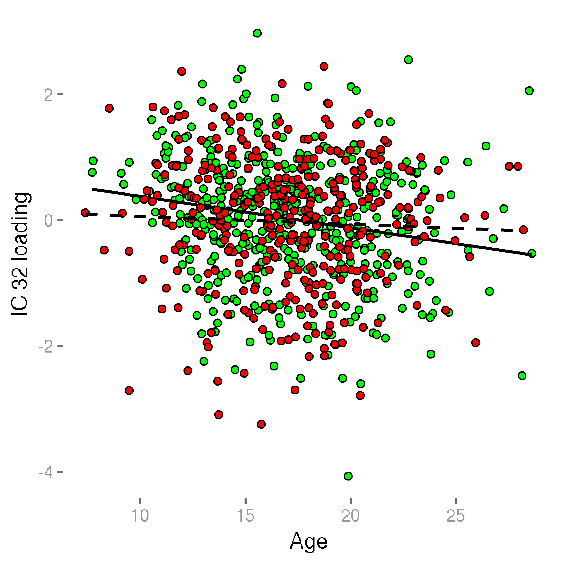

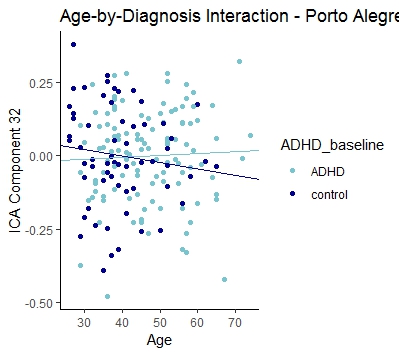

*p =* 0.272

*p =* 0.0233

*p =* 0.1794

*p =* 0.0176

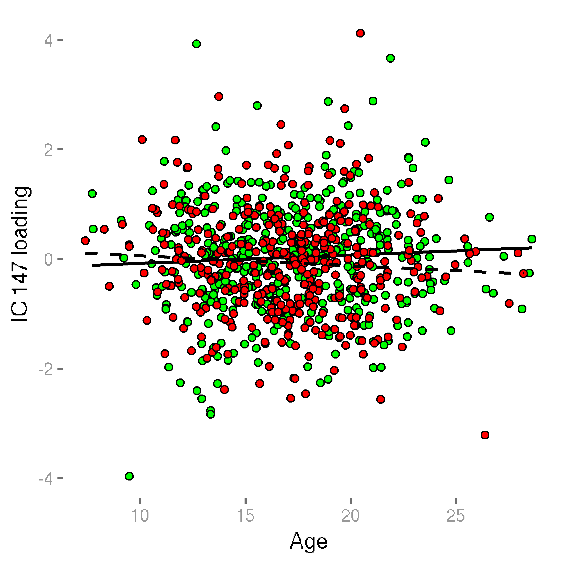

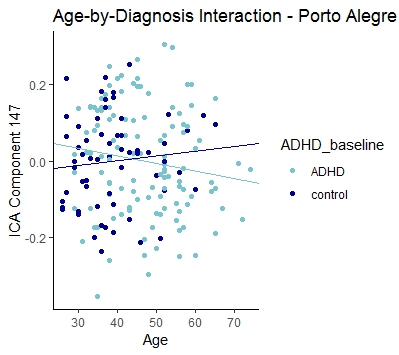

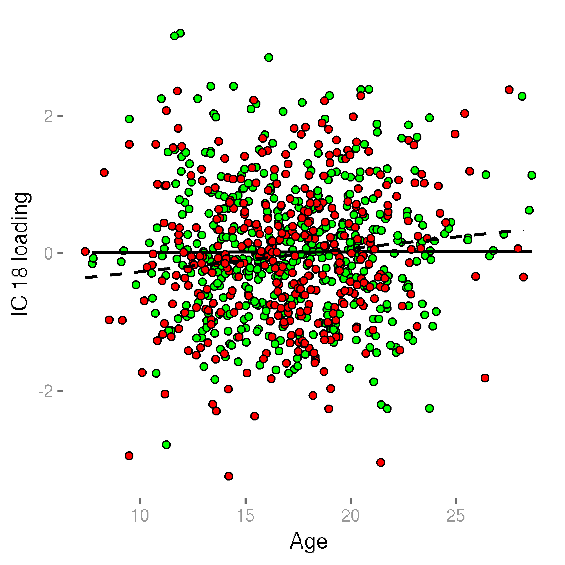

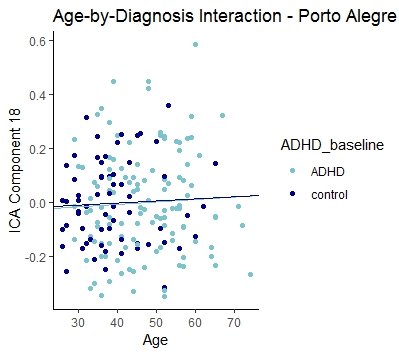

*p =* 0.0264

*p =* 0.1317

*p =* 0.0357

*p =* 0.9633

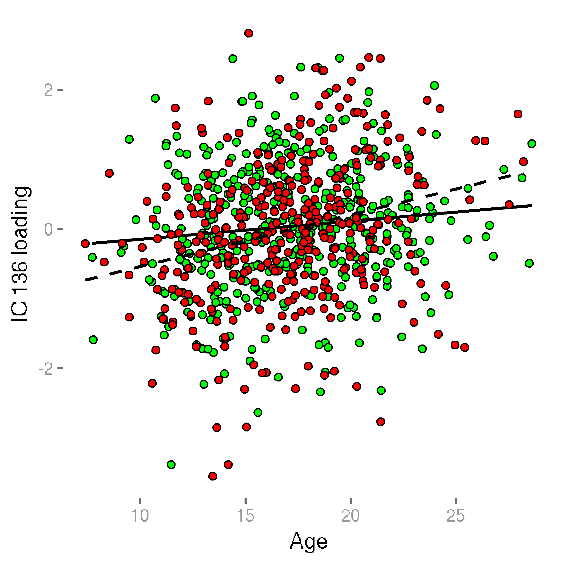

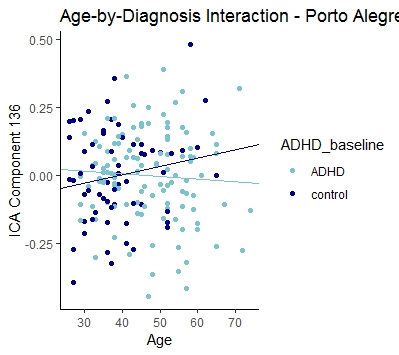

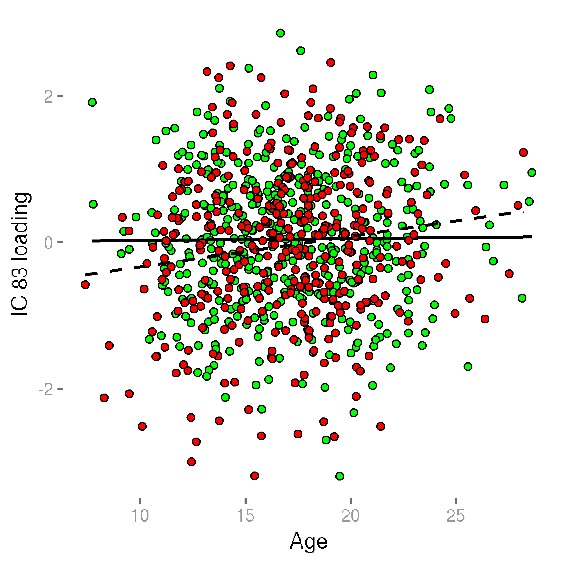

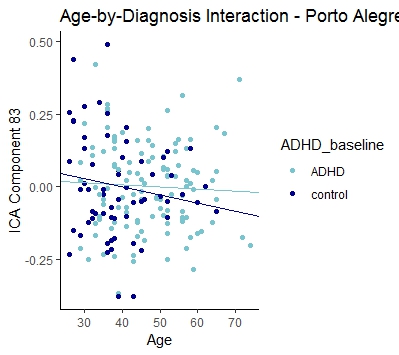

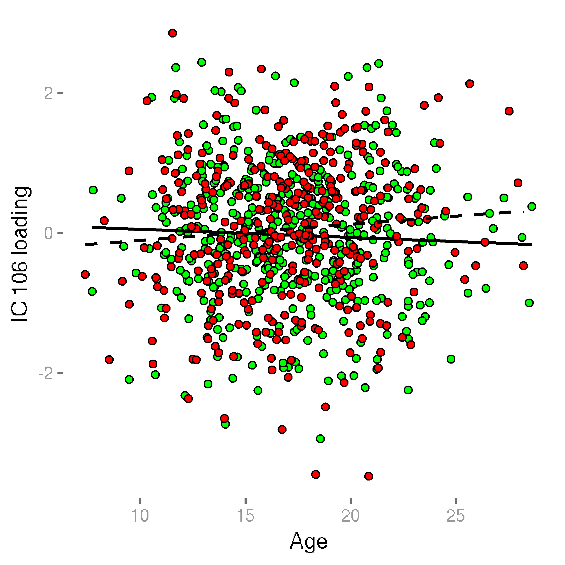

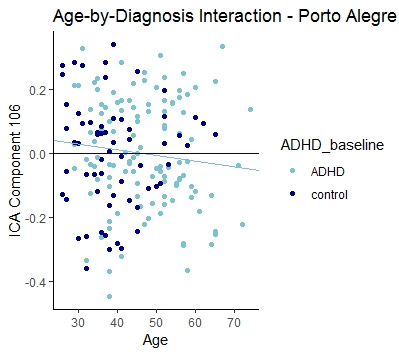

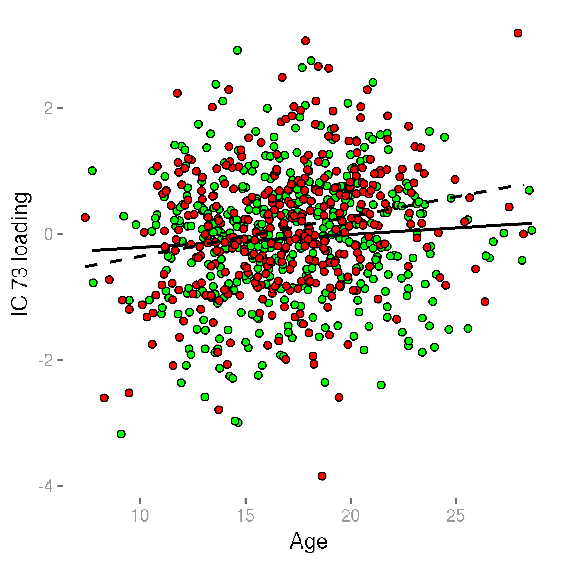

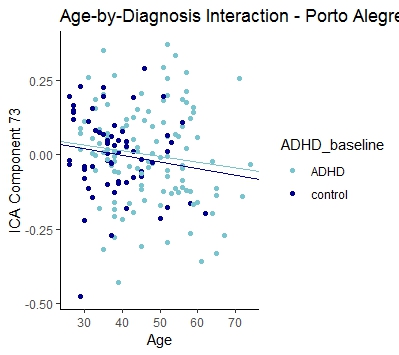

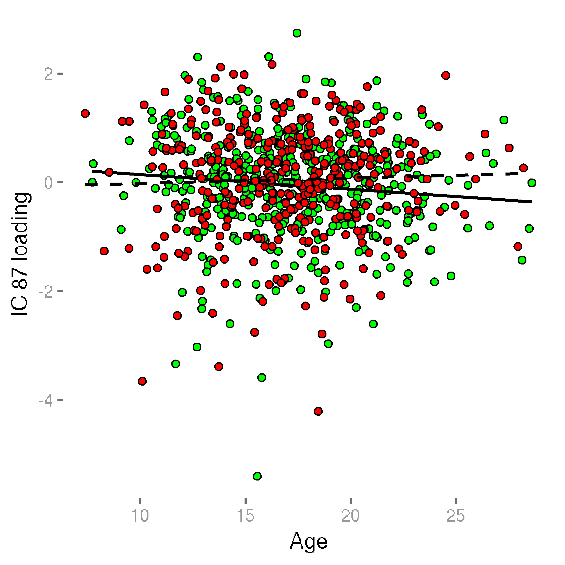

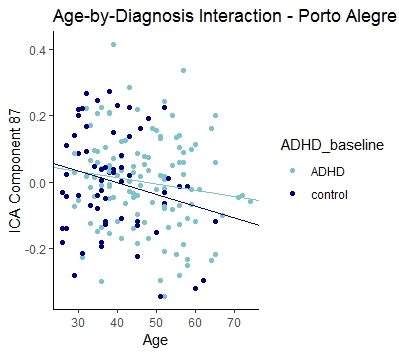

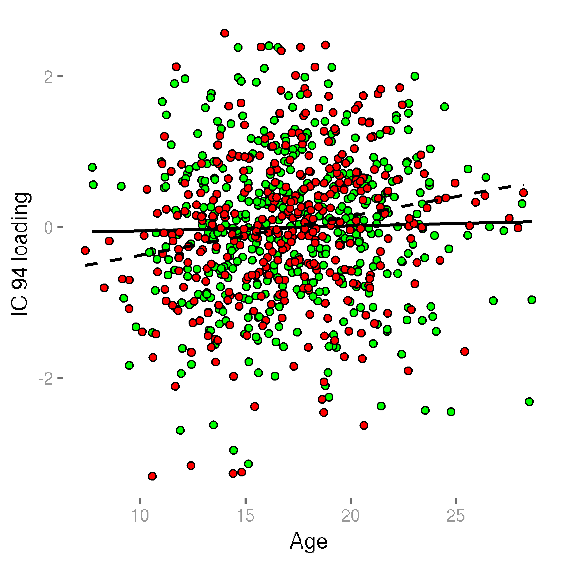

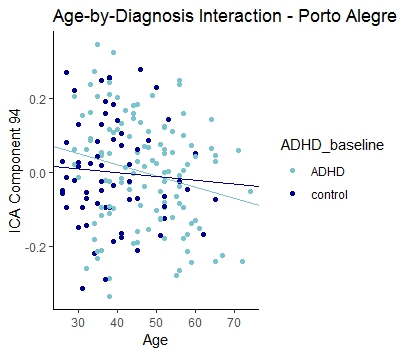

*p =* 0.902

*p =* 0.0440

*p =* 0.519

*p =* 0.0436

*p =* 0.4074

*p =* 0.0401

*p =* 0.1336

*p =* 0.0396
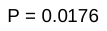

*p =* 0.0495

*p =* 0.387

*p =* 0.484

*p =* 0.0463
